# Exhaled breath condensate proteomics using amphipols improves protein detection but reveals statistical challenges in respiratory disease biomarker discovery

**DOI:** 10.1101/2025.11.21.689702

**Authors:** Eline Schillebeeckx, Andrea Argentini, Jo Raskin, Jan P. van Meerbeeck, Kris Gevaert, Kevin Lamote, Esperanza Fernández

## Abstract

**Background:** Invasive diagnostic techniques for pleural mesothelioma (PM) delay early detection, underscoring the need for non-invasive biomarkers. Exhaled breath condensate (EBC) is a promising matrix but has extremely low protein concentration. This study applies an amphipol (APol)-based concentration method to enhance detection of potential PM biomarkers in the EBC proteome.

**Methods:** Four replicate EBC samples from a healthy individual were processed using APols or lyophilization. Subsequently, 145 EBC samples from PM patients (n=25), lung cancer patients (n=28), asymptomatic asbestos-exposed individuals (n=60) or healthy controls (n=32) were concentrated with APols and analysed by liquid chromatography coupled with tandem mass spectrometry. Technical variability from batch effects and collection devices was addressed using omicsGMF, for simultaneous imputation and covariates correction. Statistical analysis employed linear models with technical and clinical covariates.

**Results:** APols outperformed lyophilization, yielding 1.7-fold higher proteome coverage and improved reproducibility. Although high data missingness was observed, proteome analysis across 145 samples identified 248 unique proteins. OmicsGMF corrected the substantial variability, while maintaining biological signal. Whereas unsupervised clustering did not reveal distinct disease-specific patterns, supervised statistical analysis detected overall subtle differences between clinical groups (PERMANOVA p-value = 0.008), though no individual proteins reached statistical significance after adjustment for technical and clinical covariates and correction for multiple testing.

**Conclusion:** APols outperforms lyophilization for concentrating proteins in low-abundant EBC samples, enabling enhanced proteome coverage for biomarker discovery studies. However, the inherent high variability of EBC proteomes has a strong effect on detecting possible biomarkers across clinical groups.

**Key message:** *What is already known on this topic:* Exhaled breath condensate (EBC) is an attractive non-invasive biofluid for respiratory disease biomarker discovery, but no protein biomarkers have reached clinical application despite extensive investigation.

*What this study adds:* Using superior APols-based protein concentration and rigorous analytical and statistical methods in 145 EBC samples, we demonstrate that fundamental limitations – 96% peptide missingness, 80% skin-protein contamination, high variability, low protein identifications and low fold-changes– make EBC unsuitable for reliable protein biomarker discovery, when proper covariate adjustment is applied.

*How this study might affect research, practice or policy:* Researchers pursuing EBC proteomics should be cautious, as inadequate handling of missingness and variability, particularly without proper covariate adjustment, may yield findings that do not reflect true biological differences.

## Introduction

Pleural mesothelioma (PM) is an aggressive cancer affecting the pleural lining surrounding the thoracic cavity. Despite recent advancements in therapy, the prognosis remains poor, with a five-year survival rate of 5-10% [1]. Biomarkers that can expedite and improve diagnosis are expected to reduce PM-related mortality. However, the only FDA-approved biomarker for PM is mesothelin measured in blood plasma, which unfortunately lacks specificity and is only approved for monitoring the treatment of epithelioid and biphasic mesothelioma. Therefore, mesothelin is not suitable for diagnostic or surveillance purposes [2]. Hence, the search for new diagnostic biomarkers and targetable molecules is ongoing [3–6]. Towards this end, exhaled breath condensate (EBC) has gained interest as a matrix of possible biomarkers as it contains a plethora of components including metabolites, proteins, DNA and RNA [7]. Its key advantage lies in the complete non-invasiveness of the sampling technique while providing molecular information about the distal airways and the composition of the airway lining fluid. Research into inflammation and oxidative stress, two main pathogenic processes of respiratory diseases, has revealed significant differences in 8-isoprostane and hydrogen peroxide levels between subjects with asbestosis and healthy controls ^12^. Furthermore, analysis of other metabolites in EBC has yielded promising results in chronic airway diseases, including cystic fibrosis, asthma and chronic obstructive pulmonary disease [7,10].

On the other hand, EBC proteomics faces significant analytical challenges, including low protein concentrations – as condensed water constitutes over 99% of EBC – which result in low numbers of identified proteins and high missing data rates, creating statistical hurdles for robust biomarker discovery [11]. To address this issue, most EBC proteome studies utilize lyophilization to concentrate proteins prior to analysis [7]. However, lyophilization is rather complex and was reported to induce protein aggregation resulting in poor protein re-solubilization which is needed for further analysis [12]. Here, we introduce an EBC protein concentration method involving the use of amphipols (APols). APols are polymers originally used for membrane protein trapping and stabilization, but that have also demonstrated utility in the broader context of proteome extraction and concentration [13]. They are compatible with trypsin digestion and do not interfere with the analysis of the resulting peptide mixtures by means of mass spectrometry. We therefore hypothesized that concentrating highly diluted protein mixtures using APols could be exploited in protein biomarker discovery pipelines.

In this study, we first compared the efficiency of APols-based protein concentration with lyophilization for EBC sample processing. Then, we evaluated the suitability of EBC for proteomic biomarker discovery by analysing individual EBC samples from pleural mesothelioma (PM) patients, lung cancer (LC) patients, asbestos-exposed individuals (AEX), and healthy controls (HC) using liquid chromatography coupled to tandem mass spectrometry (LC-MS/MS). The mass spectrometer was operated in data independent acquisition (DIA) mode to increase the overall analytical depth and improve coverage of low-abundance peptides. Through comprehensive statistical analysis with proper covariate adjustment, we aimed to identify potential protein biomarkers and to critically assess the fundamental capabilities and limitations of EBC as a matrix for respiratory disease protein biomarker discovery.

## Methods

### Study design and subject details

A cross-sectional, case-control study was conducted to recruit all participants. The study was approved by the Ethics Committee of the Antwerp University Hospital (Belgian registration number B300201837007) and conducted in accordance with the Helsinki Convention. The cohort consisted of 145 participants: 25 PM patients, 28 LC patients, 60 AEX controls and 32 HC. All participants were recruited between October 2019 and November 2021. PM and LC patients were included after referral through the Thoracic Oncology Department of the Antwerp University Hospital (Belgium). PM diagnosis was histologically confirmed and the patients were treatment-naïve at the time of inclusion. Individuals with a documented history of professional asbestos exposure, both asymptomatic and with a benign asbestos-related disease (pleural plaques, pleuritis and/or asbestosis), were recruited through the occupational health departments of three companies known for their historical involvement in asbestos processing. Healthy controls without known occupational asbestos exposure were recruited through advertisement on the website of the Antwerp University Hospital, posters and advertisement in a local magazine. Upon enrolment, all participants provided written informed consent and completed two questionnaires aimed at gathering demographic information and details about their history of asbestos exposure.

### Sample collection

EBC samples were collected using either the EcoScreen 2 (Jaeger Corporation, Germany;-20 °C) or the TurboDECCS (Medivac, Italy;-10 °C), according to the ERS guidelines [14]. Participants were instructed to fast for 2 h prior to the study visit and rinse their mouth with distilled water before sample collection [15]. During a 20-minutes period, participants were asked to breathe at tidal volume while wearing a nose clip to prevent nasal inhalation. Following collection, 10 µL /mL EDTA was added to each sample. On average, 2.26 mL of EBC was collected per participant (EcoScreen 2: 3.04 mL; TurboDECCS: 1.83 mL). The collected samples were divided into 1 mL aliquots and stored at-80 °C until analysis. Both the researcher and the participant wore gloves during the EBC sampling procedure to minimize epidermal keratin contamination. The EBC devices are equipped with either a saliva trap or a swan-neck to reduce saliva contamination. Each participant provided one EBC sample, except for one healthy control, who provided eight samples (four of 1 mL and four of 2 mL).

Method details on Sample preparation, mass spectrometry analysis and statistical analysis are provided in the Supplemental material.

## Results

### Amphipols facilitates protein identification in EBC samples

Four EBC samples from a control subject were collected and proteins were concentrated using APols-mediated precipitation (figure 1a). To assess APols efficiency, we compared the peptides and proteins identified in the APols precipitates with those remaining in the supernatant (figure 1b). Following data filtering, 1,330 peptides and 117 proteins were uniquely identified by LC-MS/MS analysis in the APols samples, with 49% and 73% of the peptides and proteins, respectively, being commonly identified across all the samples (supplemental tables 1-2). Quantitative analysis of 362 peptides identified in both APols precipitates and supernatant fractions revealed efficient APols-mediated protein concentration, with a median recovery of 83.7% (Interquartile Range: 70.2 - 91.3%) (figure 1c) (supplemental table 3). The majority of peptides demonstrated strong recovery, with 104 (28.7%) peptides showing >90% recovery efficiency (supplemental figure 1). Importantly, the cumulative intensity of peptides in the APols precipitate represented 80% ± 1.2% (mean ± SD, n=3) of the total signal (APols + SN), representing 4.0 ± 0.3-fold enrichment in the APols fraction (supplemental table 4). These results confirm that APols effectively concentrates proteins from dilute EBC samples.

**figure 1.**
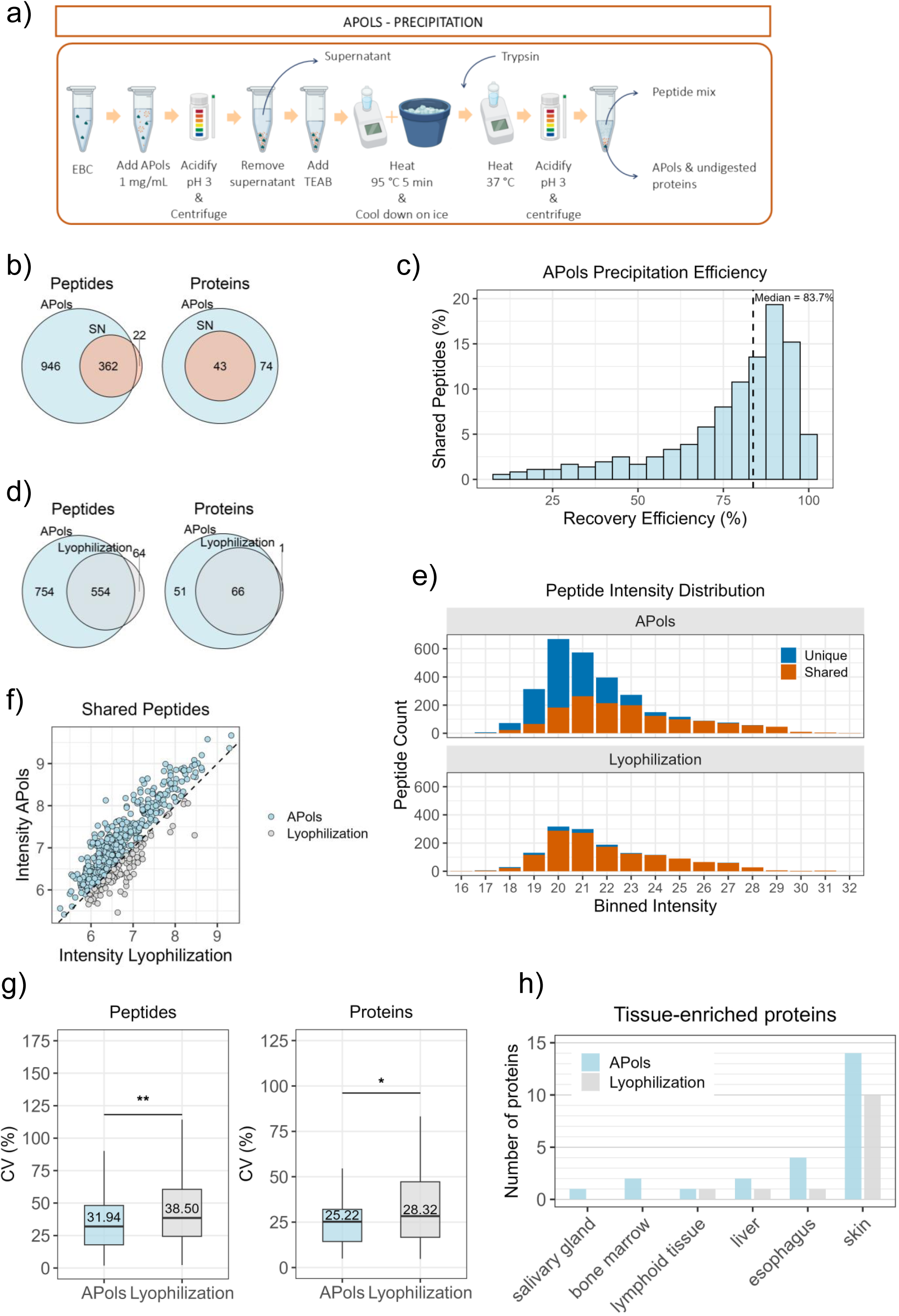
Overview of the method used and the results obtained upon optimizing the EBC proteome concentration technique. (**a**) Overview of the Apols precipitation method. **(b)** Venn diagrams showing the overlap of unique peptides and proteins identified in at least three replicates of the APols-precipitate (APols) and APols supernatant (SN). **(c)** Recovery metrics for shared peptides identified in APols precipitates and SN (n = 362). The recovery percentage was calculated as (Intensity_APols / (Intensity_APols + Intensity_SN)) × 100. **(d)** Venn diagram comparing the unique peptides and proteins identified in APols-precipitate or the lyophilized EBC samples. **(e)** Peptide intensities (log₂-transformed) were binned and plotted as stacked bars for each condition (APols and Lyophilization). Bars represent the number of peptides within each log₂ intensity bin, with each bin spanning one log_2_ unit. Shared peptides are shown in orange, unique peptides in blue. **(f)** Distribution of shared peptide intensities calculated as the mean of intensities across replicates of both groups and represented as log_10_ of the mean (n = 554). (**g**) Distribution of the coefficient of variation (CV) of the normalized raw intensities, and the label free quantification (LFQ) intensities of the shared peptides (n = 253) and proteins (n = 33), respectively, identified in at least three replicates of the APols and lyophilization methods. Paired Wilcoxon signed-rank test, peptides *p*-value = 0.002; proteins *p*-value = 0.035. **(h)** Histogram indicating the number of proteins annotated by the Human Protein Atlas v24.0 as enriched in the indicated tissues. *APols: amphipols, EBC: exhaled breath condensate, SN: supernatant.* * *p* < 0.05, ** *p* < 0.01, by the statistical test indicated.

Next, we compared the efficiency of APols-mediated concentration to lyophilization, a commonly used method for concentrating dilute biofluids [16]. Following lyophilization, a total of 618 peptides and 67 unique proteins were identified, of which 90% of the peptides and 98% of the proteins were also identified in the APols precipitates (figure 1d) (supplemental tables 1-2). Keratin KRT6C, expressed in squamous epithelium, was the sole protein only identified upon lyophilization. In contrast, APols identified 754 unique peptides and 51 unique proteins not detected by lyophilization, representing a substantial increase in proteome coverage (figure 1d).

Binned intensity analysis revealed that APols enabled the identification of substantially more unique peptides than lyophilization, particularly in the mid-to-high intensity range (intensity bins 19-23) (figure 1e). Shared peptides between methods (n = 554), exhibited similar intensity distribution patterns; however, APols consistently yielded higher signal intensities, as evidenced by 81.2% of peptides plotting above the line of equal intensity in the correlation analysis with a median of 2.16-fold higher levels in APols (paired Wilcoxon signed-rank test, *p* < 0.001) (figure 1f, supplemental figure 2).

Since protein extraction is a major source of experimental variability [17], we assessed the quantification precision of both methods by measuring the CV of shared peptides and proteins identified in at least three replicates in each group. For the 253 shared peptides across replicates, APols precipitation demonstrated significantly improved peptide-level quantification reproducibility (median CV: 31.94%) compared to lyophilization (median CV: 38.50%, paired Wilcoxon signed-rank test, *p* < 0.01) (figure 1g). Similarly, for the 33 shared proteins quantified in both methods across replicates, APols showed significantly lower median CV (25.22%) compared to lyophilization (28.32%, paired Wilcoxon signed-rank test, *p* < 0.05) (figure 1g). These findings indicate that APols enhances quantification precision when compared with lyophilization.

Annotation of tissue expression profiles using the Human Protein Atlas [18] revealed that 24 proteins were annotated as tissue-enriched across both methods (figure 1h, supplemental table 5). Skin was the most prevalent annotated tissue, with 14 proteins classified as skin-enriched, consistent with the expected contribution of skin to the EBC proteome [19]. Four proteins (KRT4, TGM1, TGM3 and KRT78) annotated as esophagus-enriched, were identified among the APols-precipitated proteins, while only KRT78 was identified in lyophilized samples (figure 1h).

Altogether, these results show that APols polymers are highly effective for concentrating proteins from low-abundance matrices, yield superior signal intensities and quantification precision compared with lyophilization, and enhance both proteome coverage and the detection of tissue-specific proteins from the respiratory tract.

### Identification of candidate diagnostic biomarkers for PM

We applied the APols-driven precipitation method coupled to MS-based proteomics to perform proteome profiling of 145 EBC samples. The patient characteristics are shown in table 1. Overall, more male participants were included in the study and PM and LC patients were significantly older than the control groups. Following data filtering, a total of 10,493 peptide precursors corresponding to 8,437 unique peptide sequences were identified across 145 samples. Precursor intensities did not show any significant difference among the samples collected with TurboECCS and EcoScreen devices (figure 2a).

**figure 2.**
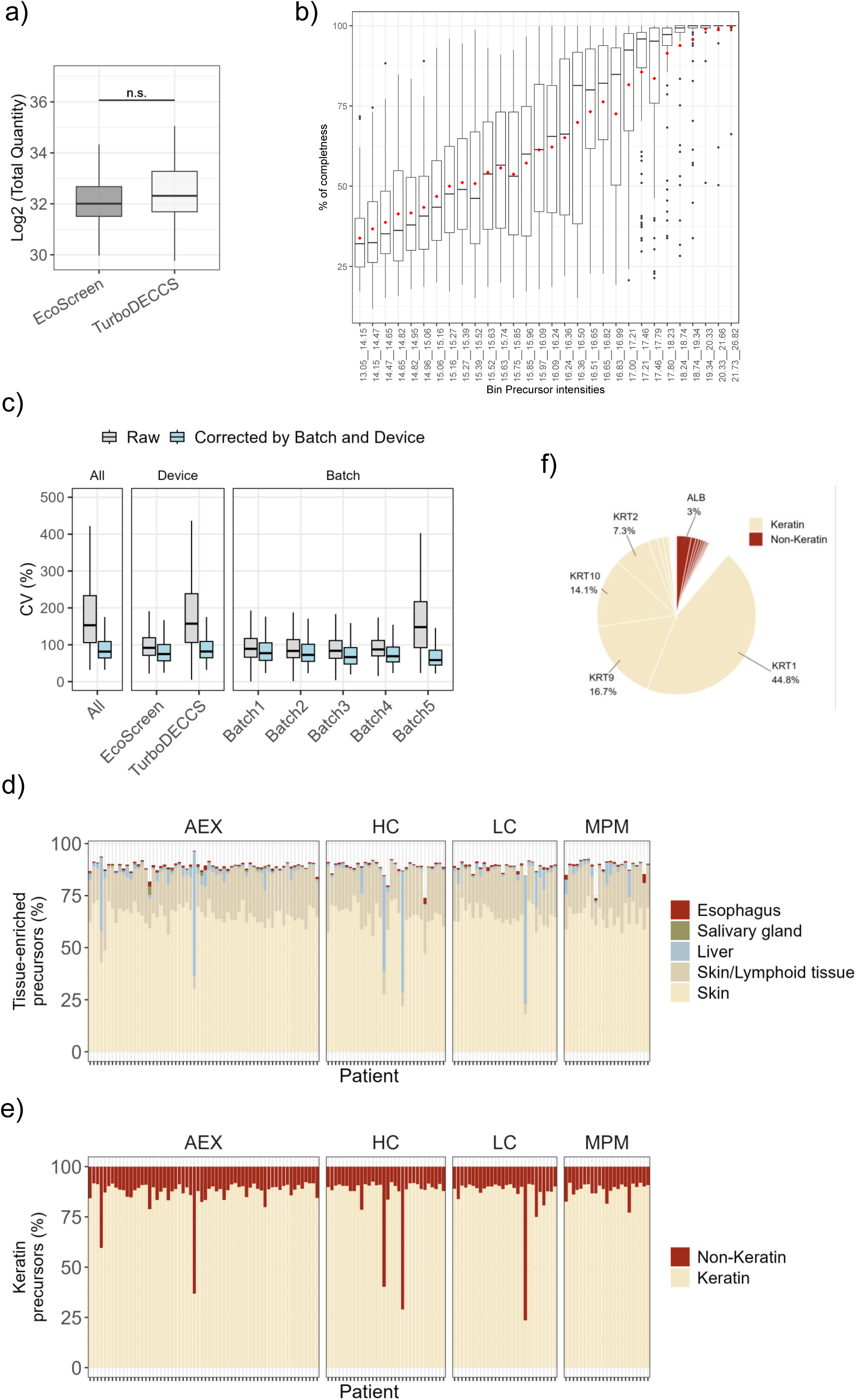
Characterization of EBC proteome composition, data quality and technical variability. **(a)** Precursor intensity distributions of the samples across the two EBC collection devices used: EcoScreen and TurboDECCS. **(b)** Distribution of the intensities of 2,374 precursors (x-axis) plotted against the percentage of completeness (y-axis). Similar number of precursors are binned (n = 79-80). Intensities are normalized and log_2_ transformed. Boxes show the interquartile range and the median. Red dots show the averages of the completeness percentage. **(c)** Box plots showing the coefficient of variation (CV) distribution for 2,374 precursors measured across different sample groupings before (raw, in grey) and after batch and device correction with omicsGMF (in blue). Panels indicate the CV of the precursors in all the samples (All) and grouped by Device and Batch, as indicated. Each box represents the interquartile range (25^th^-75^th^ percentile), with the median indicated by the central line. Whiskers extend to the most extreme data points within 1.5 times the interquartile range. Raw CVs (%): All precursors = 157.44; EcoScreen = 92.00; TurboDECCS = 161.75; Batch 1 = 89.12; Batch 2 = 83.35; Batch 3 = 83.74; Batch 4 = 87.44; Batch 5 = 147.87. OmicsGMF corrected CVs (%): All precursors = 82.86; EcoScreen = 75.03; TurboDECCS = 83.11; Batch 1 = 77.25; Batch 2 = 72.67; Batch 3 = 66.57; Batch 4 = 69.41; Batch 5 = 58.50. **(d)** Bar chart showing the percentage distribution of the 2,374 unique precursor intensities mapped to tissue-enriched proteins according to Human Protein Atlas version 24.0. Tissues enriched in more than 1% of the precursor intensities are indicated. **(e)** Bar chart showing the percentage of precursor intensities to illustrate the high abundance of keratins. **(f)** Pie chart displaying the relative spectral intensity contributions of the most five abundant proteins, Keratin-1, Keratin-2, Keratin-9, Keratin-10 and Albumin.

**Table 1.**
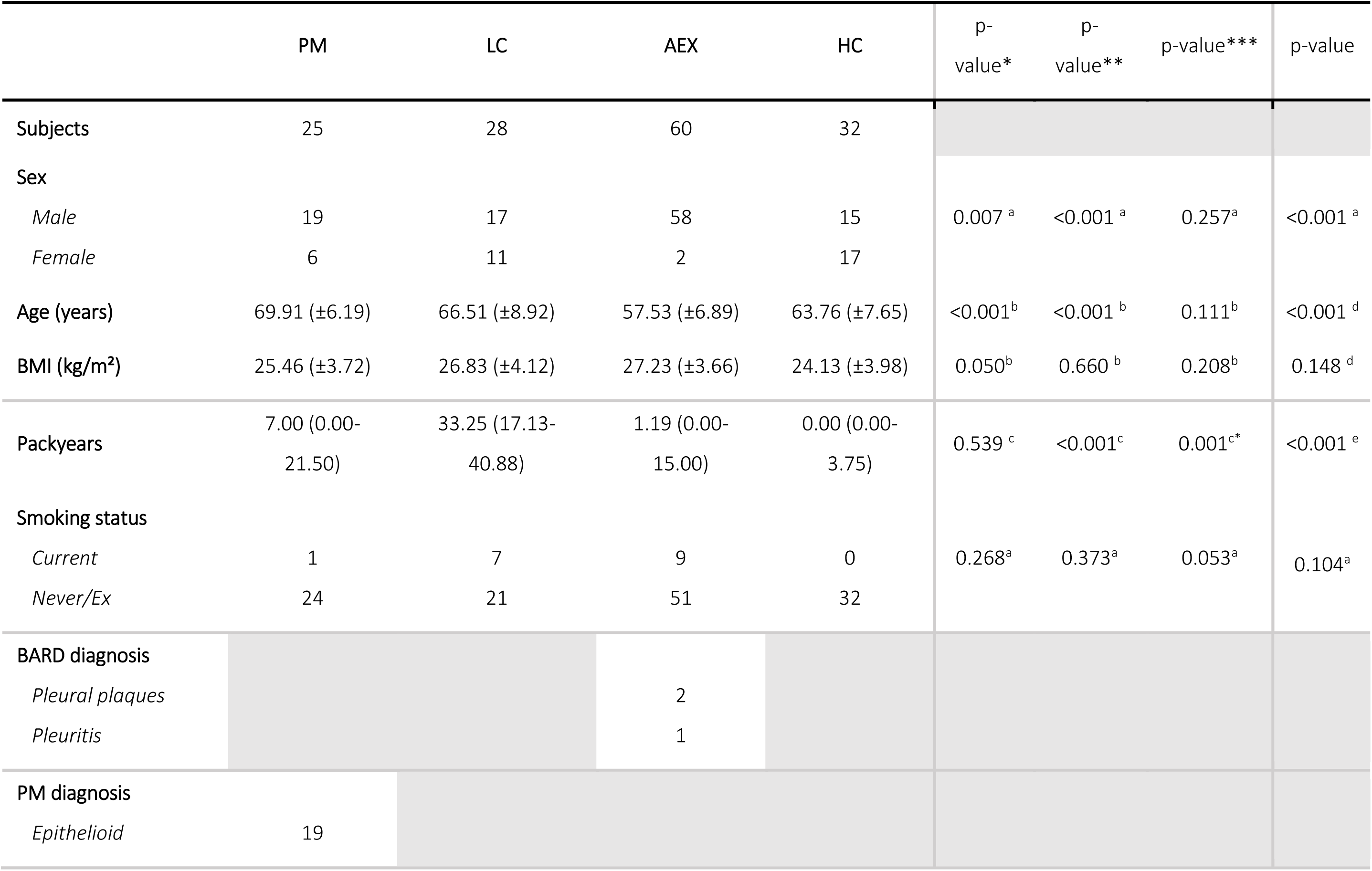
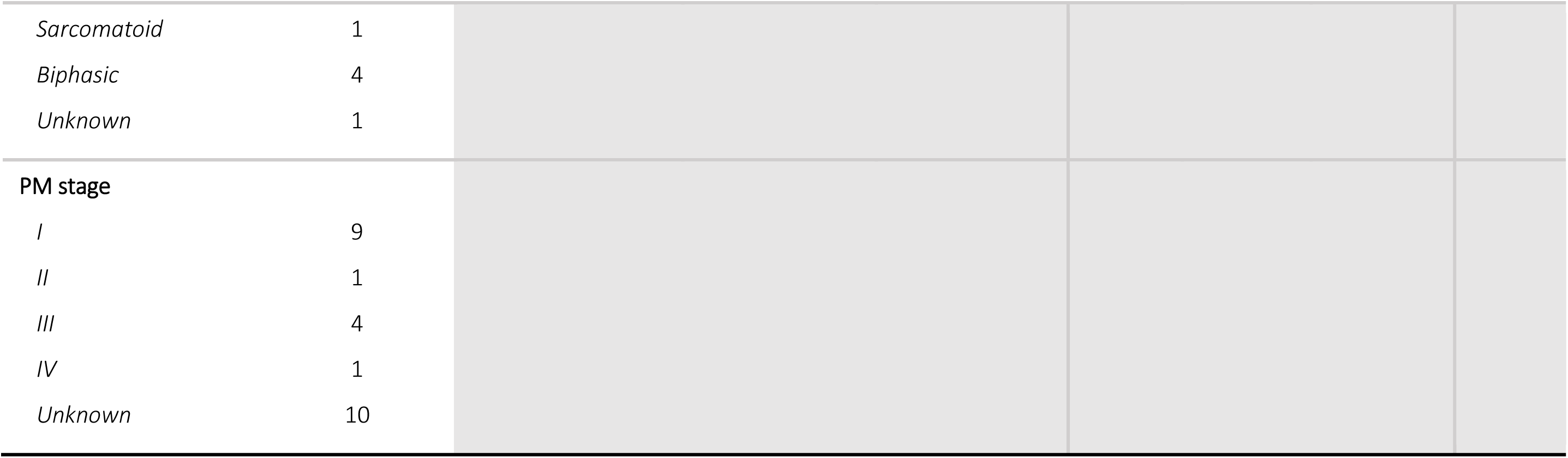
Patient characteristics. Values are presented as n, mean ±SD or median (Q1-Q3). AEX: professionally asbestos-exposed individual, HC: healthy control without known asbestos exposure, LC: lung cancer, PM: pleural mesothelioma. a Fisher’s exact test; b t-test, c Mann-Whitney U test, d One-Way ANOVA, e Kruskal-Wallis test, * comparison PM vs AEX, ** comparison LC vs AEX, *** comparison PM vs LC

|  | PM | LC | AEX | HC | p-value* | p-value** | p-value*** | p-value |
| --- | --- | --- | --- | --- | --- | --- | --- | --- |
| **Subjects** | 25 | 28 | 60 | 32 |  |  |  |  |
| **Sex** |  |  |  |  |  |  |  |  |
| *Male* | 19 | 17 | 58 | 15 | 0.007<sup>a</sup> | <0.001<sup>a</sup> | 0.257<sup>a</sup> | <0.001<sup>a</sup> |
| *Female* | 6 | 11 | 2 | 17 |  |  |  |  |
| **Age (years)** | 69.91 (±6.19) | 66.51 (±8.92) | 57.53 (±6.89) | 63.76 (±7.65) | <0.001<sup>b</sup> | <0.001<sup>b</sup> | 0.111<sup>b</sup> | <0.001<sup>d</sup> |
| **BMI (kg/m<sup>2</sup>)** | 25.46 (±3.72) | 26.83 (±4.12) | 27.23 (±3.66) | 24.13 (±3.98) | 0.050<sup>b</sup> | 0.660<sup>b</sup> | 0.208<sup>b</sup> | 0.148<sup>d</sup> |
| **Packyears** | 7.00 (0.00-21.50) | 33.25 (17.13-40.88) | 1.19 (0.00-15.00) | 0.00 (0.00-3.75) | 0.539<sup>c</sup> | <0.001<sup>c</sup> | 0.001<sup>c*</sup> | <0.001<sup>e</sup> |
| **Smoking status** |  |  |  |  |  |  |  |  |
| *Current* | 1 | 7 | 9 | 0 | 0.268<sup>a</sup> | 0.373<sup>a</sup> | 0.053<sup>a</sup> | 0.104<sup>a</sup> |
| *Never/Ex* | 24 | 21 | 51 | 32 |  |  |  |  |
| **BARD diagnosis** |  |  |  |  |  |  |  |  |
| *Pleural plaques* |  |  | 2 |  |  |  |  |  |
| *Pleuritis* |  |  | 1 |  |  |  |  |  |
| **PM diagnosis** |  |  |  |  |  |  |  |  |
| *Epithelioid* | 19 |  |  |  |  |  |  |  |

**Table 1:** Patient characteristics. Values are presented as n, mean ± SD or median (Q1-Q3).
|  |  |
| --- | --- |
| *Sarcomatoid* | 1 |
| *Biphasic* | 4 |
| *Unknown* | 1 |
| **PM stage** |  |
| *I* | 9 |
| *II* | 1 |
| *III* | 4 |
| *IV* | 1 |
| *Unknown* | 10 |

Missing values in MS-based protein biomarker discovery remain a challenge, particularly in liquid biopsies, where protein concentrations vary widely or protein abundance is very low, as in EBC samples [20]. Peptides exhibited high missingness: 2,374 precursors were identified in at least 30% of samples within one clinical group, while only 346 peptides – 14.6% of the filtered dataset – were identified across all samples (supplemental figure 3). Peptide intensity strongly correlated with completeness (Spearman r = 0.991, *p*-value < 0.001), with high abundance peptides showing higher completeness across samples (figure 2b). This relationship remained consistent when samples collected by different EBC devices were analysed separately (supplemental figure 4), demonstrating that the intensity-completeness correlation is device-independent. These results indicate that missingness is predominantly driven by low protein abundance rather than technical-related variation.

Next, we sought to assess the variability of the 2,374 precursors across samples. The CV revealed substantial variability at sample level with a median CV of 157.4% across all samples (figure 2c). Variability across experimental batches was comparable, with the exception of batch 5, which exhibited higher variability than batches 1-4. In addition, a difference in variability was observed between the two collection devices used, suggesting that device-specific technical variation may contribute to overall data heterogeneity (figure 2c). To address this, we applied omicsGMF, a matrix factorization method that integrates imputation and factor correction into a unified framework [21]. This method has been shown to effectively reduce dimensionality and facilitates the identification of differentially expressed proteins in datasets affected by strong batch effects and high levels of missing data [21]. Following correction for batch and device effects, the distribution of CVs across samples became more uniform. Nevertheless, the overall CV remained elevated, ranging between 57% and 83%, indicating additional sources of variability persisted beyond the corrected factors (figure 2c). Similar variation was observed among precursors identified within clinical groups, further supporting sample heterogeneity rather than clinical variability (supplemental figure 5).

Next, we investigated the tissue origin of the 2,374 precursors using the RNA and protein tissue classification from the Human Protein Atlas [18]. Only precursors mapped to a protein annotated as enriched in one tissue (tissue-enriched) were retained. On average, 70% of the total spectral intensity mapped to proteins enriched in the skin, 17% in the skin/lymphoid tissue and less than 2% were annotated as enriched in the esophagus and squamous epithelium-tissues, indicative of the upper-respiratory tract (figure 2d) (supplemental table 6). The annotation of skin as the most prevalent tissue is due to the presence of keratins (figure 2e) [19]. Keratins-1,-2 and-10 are strongly expressed in the suprabasal layers of epidermis and Keratin-9 is enriched in the palmoplantar skin, and peptides from both cover about 80% of the total spectral intensity (figure 2f).

Precursor intensities corrected by device and batch effects were summarized into proteins yielding a total of 248 unique proteins across all the samples (supplemental table 7). Global protein abundance was comparable across clinical groups, with no significant differences between AEX, HC, LC and PM (One-way ANOVA, *F* (3, 141) = 1.26, *p* = 0.29) (figure 3a). To assess protein quantification variability within each clinical group, we calculated the CV for individual proteins. Overall, 38.7%, 29.8%, 49.6%, and 35.4% of the proteins quantified across the AEX, HC, LC and PM groups, respectively, had CVs below 50% (figure 3b). This indicates that the majority of proteins (>50% in most groups) exhibited high variability within clinical groups, with CVs exceeding 50%.

**figure 3.**
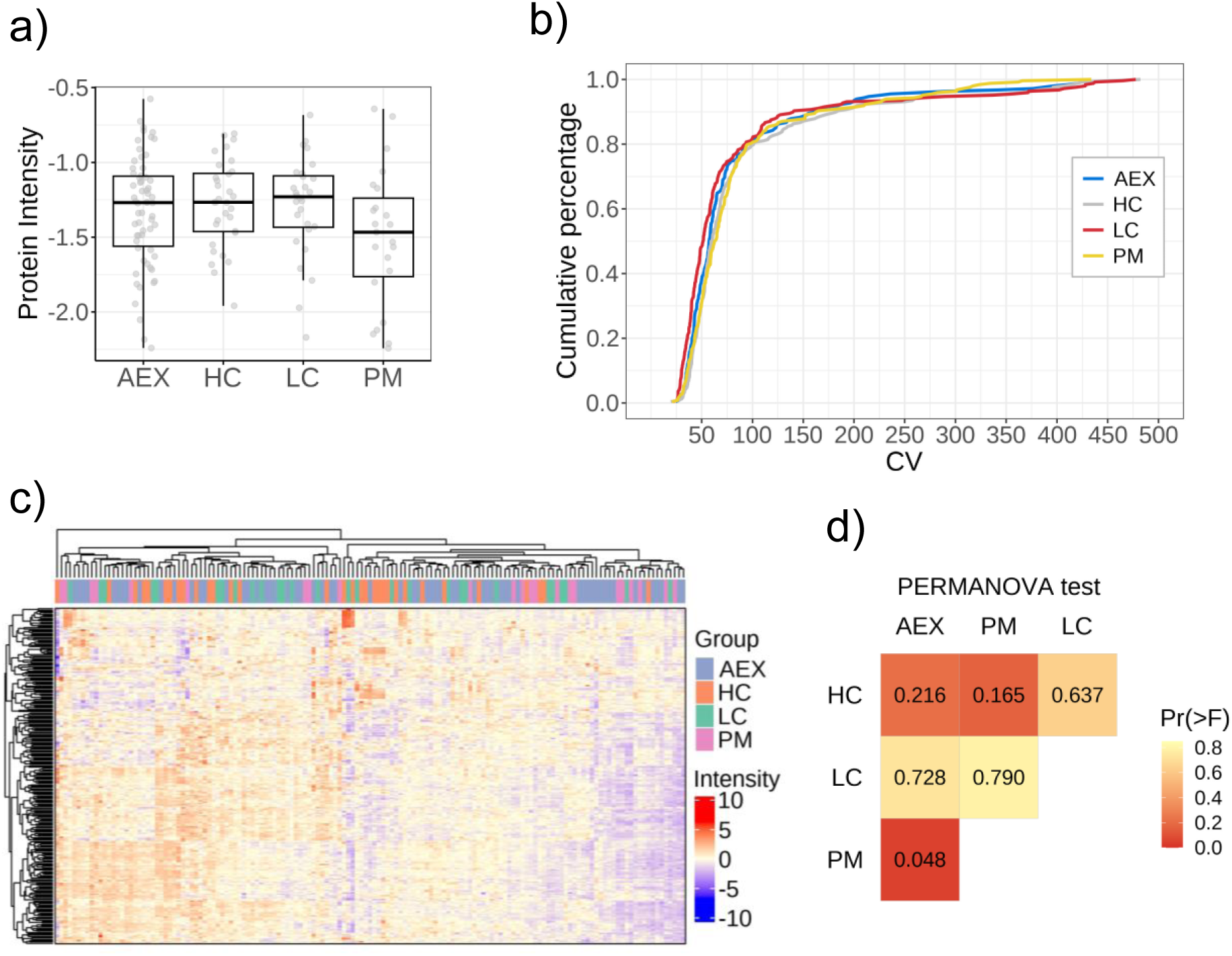
Protein expression analysis reveals comparable global abundance but subtle molecular differences between clinical groups. **(a)** Box plot showing the distribution of normalized protein abundance across the four clinical groups. One-way ANOVA, F(3, 141) = 1.26, p = 0.29. Each box represents the interquartile range with the median indicated by the central line. **(b)** Cumulative distribution plot showing the cumulative percentage of proteins (y-axis) plotted against the coefficient of variation (CV) values (x-axis) for each clinical group. **(c)** Dendrogram of the hierarchical clustering of 145 EBC samples based on 248 protein intensities. Clinical groups (AEX, HC, LC, PM) are color-coded as indicated. Protein intensities are log_2_ transformed and z-score normalized. **(d)** Heatmap of the results of pairwise Permutational Multivariate Analysis of Variance (PERMANOVA) comparison tests between clinical groups, indicating the p-values (Pr(>F)). AEX: asbestos-exposed, HC: healthy controls, LC: lung cancer, and PM: malignant pleural mesothelioma.

**figure 4:**
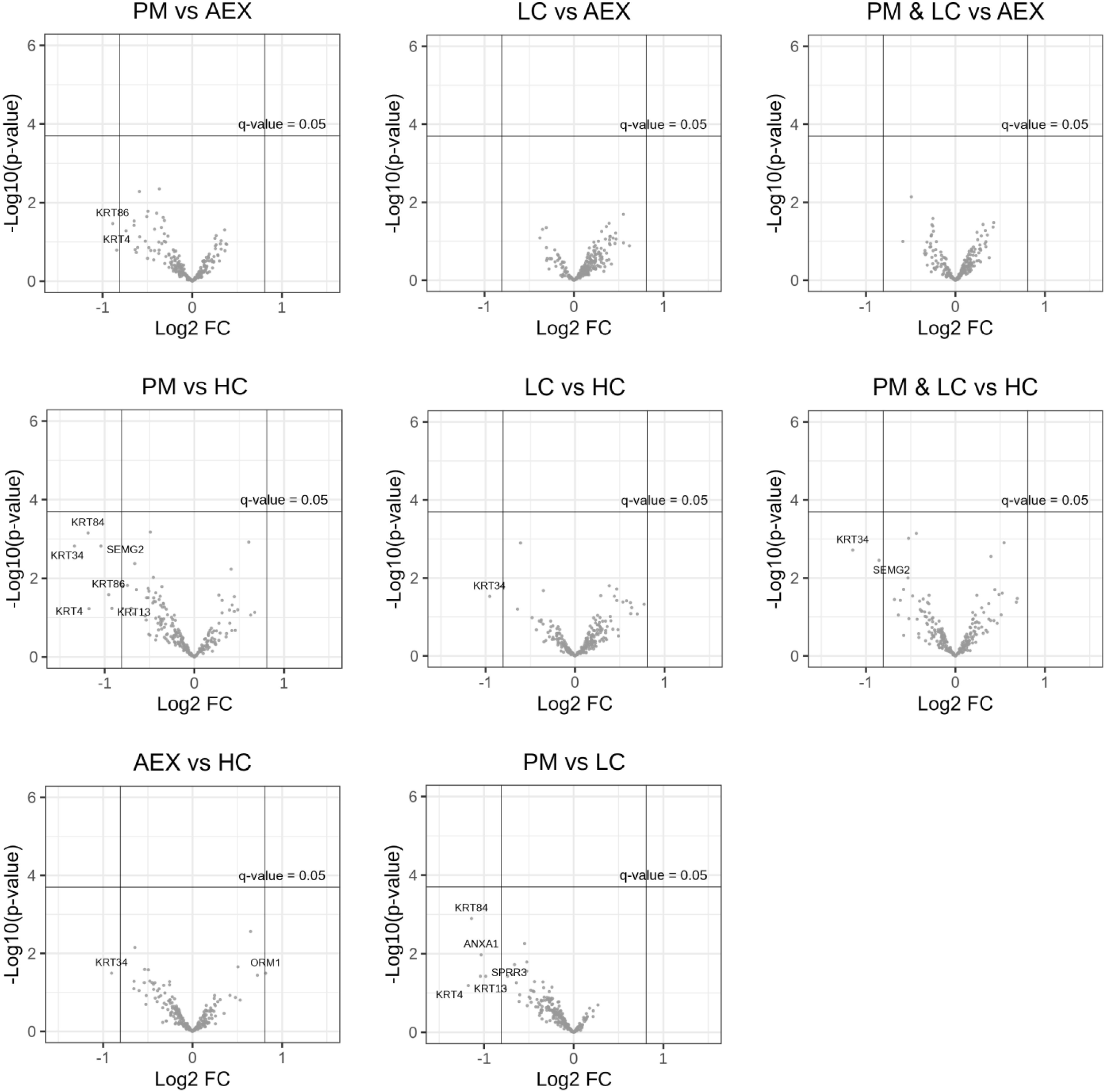
Differential expression analysis across clinical groups. Volcano plots representing the results of the statistical linear model analysis between the indicated contrasts of the clinical groups. Significance threshold q-value (adjusted p-value) < 0.05. Gene names corresponding to proteins with 1.75-fold change (log_2_ fold-change = 0.8073) are indicated.

To evaluate the effect of potential technical covariates, we performed a Principal Component Analysis (PCA) on the protein intensity data after correction for device and batch effects. This analysis revealed no separation of samples by device type or processing batch along the first two principal components, suggesting that these technical variables did not drive major sources of variation in the dataset (supplemental figure 6). This supports the effectiveness of the correction procedure and indicates that downstream analyses are unlikely to be confounded by batch or device artefacts.

To assess whether the four sample cohorts exhibited distinct protein expression signatures, we performed both unsupervised hierarchical clustering analysis (HCA) and supervised statistical testing across the 248 proteins measured in all EBC samples. HCA did not reveal clear group-specific clusters, as evidenced by the scattered distribution of clinical groups throughout the protein dendrogram (figure 3c). Quantitative analysis supported this observation: the average silhouette width was 0.17, indicating weak clustering structure (values >0.25 suggest meaningful clusters), and the Adjusted Rand Index (ARI) comparing clinical grouping to molecular clusters was 0.002, indicating no correspondence between protein clustering patterns and clinical classifications.

However, supervised statistical analysis revealed subtle, yet statistically significant molecular differences between clinical groups. A PERMANOVA test identified significant group-level differences (R² = 0.037, *p* = 0.008), although the effect size was small, with clinical groupings accounting for only 3.76% of total variance. Pairwise PERMANOVA comparisons showed significant differences between the AEX and PM groups (F = 1.90, R² = 0.022, *p* = 0.048) (figure 3d) (supplemental table 8). These findings were supported by Analysis of Similarities (ANOSIM), which yielded a small but significant variation (Global R = 0.086, *p* = 0.007) indicating minimal between-group differences relative to within-group variation. Overall, while the unsupervised clustering did not separate sample groups, the supervised analyses suggest subtle, yet statistically significant differences in protein expression patterns, particularly between specific clinical groups.

### Differential protein expression analysis does not reveal disease-specific signatures

To identify proteins that differ significantly between clinical cohorts, we performed pairwise linear model comparisons across all group combinations, including a combined comparison of both cancer groups against the control groups. Given the subtle overall differences detected by PERMANOVA and the high within-group variability observed in the EBC proteome, we applied stringent statistical criteria and included all covariates (device, batch, age, sex, BMI, packyears and smoking status) to identify robust biomarker candidates. Even though 56 proteins showed a *p*-value < 0.05 in at least one of the contrasts, no protein remained significantly differentially expressed between any pair of clinical groups after correction for multiple testing (q-value < 0.05) (supplemental tables 8-16).

Despite the absence of statistically significant changes, we observed several proteins with moderate fold changes (> 1.75-fold change) that may be worth further investigation. In the PM vs HC comparison, epithelial keratins KRT4, KRT13, KRT34, KRT84, and KRT86 showed trends toward downregulation in PM patients, along with SEMG2. These keratins are markers of stratified squamous epithelium found in the oral cavity and esophagus (KRT4, KRT13) or hair follicles and skin (KRT34, KRT84, KRT86) [18]. The PM vs LC comparison revealed similar patterns with KRT4, KRT13, and KRT84 trending downward in PM, alongside ANXA1, an anti-inflammatory protein expressed in epithelial cells, and SPRR3, a squamous epithelial differentiation marker, downregulated in the transition from normal to high-grade intraepithelial neoplasia [18,22]. In the AEX vs HC comparison, KRT34 and ORM1 (an acute phase inflammatory protein) showed trends toward altered expression. The consistent trend of reduced epithelial keratins in PM patients across multiple comparisons may reflect mechanical and breathing pattern changes, epithelial atrophy or damage, reduced epithelial desquamation, or systemic effects of the disease, though these observations require further validation.

## Discussion

Our study demonstrates that amphipols (APols) represents a significant advancement in EBC protein concentration compared to traditional lyophilization. The superior performance of APols is evidenced by increased proteome coverage and improved signal intensity. APols enabled the identification of 2.1-fold and 1.7-fold more peptides and proteins, respectively, compared to lyophilized samples. Also, 81% of peptides commonly identified in both methods showed enhanced detection in APols. This enhancement was accompanied by improved quantification precision with lower coefficients of variation (CVs) at both peptide and protein levels. The complete protein recovery and the identification of esophagus-enriched annotated proteins such as KRT4, TGM1 and TGM3, which were not detected in lyophilized samples, highlights the enhanced sensitivity of this approach and validates the advantages of APols for proteomes with very low protein levels. Applied to a cohort of 145 EBC samples from individuals carrying pleural mesothelioma (PM), lung cancer (LC), occupationally exposed to asbestos (AEX) or healthy controls (HC), we found that after correcting for multiple covariates using omicsGMF, the high and innate variability in EBC samples did not allow for the identification of proteins that might discriminate EBC samples of patients from those of healthy individuals.

EBC has gained attention as a non-invasive source of biomarkers for the detection of respiratory diseases [7]. However, the major constrains of EBC proteomics is the low concentration of proteins, often near or under the analytical detection limit and the inherent intra-and inter-individual variability of protein concentration [15,20].

First, protein concentration methods lead to protein loss and irreversible protein aggregation, thereby introducing sample variability that has a detrimental impact on quantitative proteomics. The capacity and efficiency of APols for concentrating proteins have already been demonstrated to be superior to ultrafiltration and acid-based precipitation methods for low abundant samples, such as the cell secretome [13]. This better performance likely stems from its mechanism of action: amphipathic polymers stabilize hydrophobic protein regions while maintaining protein solubility, preventing protein aggregation and precipitation losses.

Secondly, the EBC protein content is influenced by multiple non-clinical factors that contribute to inherent sample variability, independent of disease status or analytical methodology. Bloemen *et al*. reported an inter-individual variability of 36.64% in EBC protein concentration, with age and height accounting for 39% of the total variance [20]. Smoking status has been reported to increase protein concentration up to five-fold ^29^.

Thirdly, EBC proteomics results in a high proportion of one-peptide-based protein identifications and therefore in poor overlap between samples. The majority of EBC proteome studies have identified just a few dozen of proteins [11]. The overall sensitivity for protein identification can however be improved by a data independent acquisition (DIA) MS approach, which allows the acquisition of all fragment ions, independently of the intensity of their peptide precursor ions, resulting in a better coverage of low abundant peptides [24]. Despite this improvement in analytical coverage, challenges remain in EBC proteomics as demonstrated by us and in other studies. Ma *et al*. reported the identification of 1,151 proteins in 20 EBC samples (10 healthy controls and 10 lung cancer) using a DIA-MS approach [25]. However, even with the enhanced peptide coverage provided by DIA, the inherent high missingness between EBC samples led to 72% of all identified proteins being excluded from statistical analysis due to insufficient data quality [25]. This highlights that while DIA improves peptide detection, the fundamental challenges of EBC proteomics, high inherent biological variability and low protein concentrations, persist.

Our comprehensive analysis of 2,374 precursors across 145 samples revealed the pervasive challenge of missing data in EBC proteomics, with only 14.5% of precursors identified in all samples. The strong correlation between peptide abundance and data completeness (Spearman r = 0.991) confirms that missingness is primarily driven by the inherently low protein concentrations in EBC rather than technical artefacts. This suggests that missing data patterns in EBC can be biologically meaningful rather than random, requiring specialized analytical approaches [26]. In addition to missingness, other covariates may affect the level of significance. In our study two different EBC collection devices were used. Unfortunately, during the course of the study a shift in the EBC collection approach was necessary due to the COVID-19 pandemic. As a safety precaution, we needed to shift from the EcoScreen (with reusable components) to the TurboDECCS (single-use components). As both equipment and temperature are listed amongst the main influencing factors, this could have introduced heterogeneity in the samples [7,15,27]. However, PCA analysis following device and batch correction did not show any clear clustering of the samples per device (supplemental figure 6).

To address missing data and technical variability, we applied omicsGMF, which unifies imputation and batch correction in a single framework [21], rather than performing these steps sequentially [28,29]. This approach effectively reduced the coefficient of variation to 57-83%, representing substantial improvement over uncorrected data, though the persistent high variability suggests much of the variation in EBC reflects genuine biological noise rather than correctable batch effects - which is an important finding about the nature of the EBC proteome itself. The inclusion of multiple covariates (age, sex, smoking, BMI, pack years, batch, device) is essential for valid inference but increases model complexity and reduces statistical power, a critical trade-off in samples with very low protein concentrations like EBC samples. Unlike most EBC proteome studies that use simple statistical approaches with limited covariate adjustment [19,25,30,31], our rigorous approach with 145 samples and multiple covariates created limited degrees of freedom for error estimation. Combined with high inherent variability and low numbers of identified proteins, this created a statistically challenging scenario where true biological differences may not reach significance after proper multiple testing correction. This raises a fundamental question: whether significant discoveries from simpler statistical methods using EBC proteomes represent true biological differences or statistical artefacts.

The predominance of skin-derived proteins in our EBC samples, accounting for 80% of spectral precursor intensity and consisting primarily of keratins, reflects the anatomical sampling pathway and aligns with previous findings [19,30,32]. Mucilli *et al*. demonstrated, using emPAI values, that human skin keratins constitute the major protein component in EBC, with approximately 50% of identified proteins being keratins [19]. The minimal representation of esophagus and squamous epithelium proteins (<2%) in our dataset, further supports that EBC primarily captures proteins from the upper respiratory tract and oral cavity rather than deeper lung tissues. The keratin-dominated composition of EBC, combined with the low abundance of potentially disease-relevant epithelial proteins, further limits the sensitivity to identify meaningful protein biomarkers.

In our dataset, despite the absence of statistically significant differentially expressed proteins, squamous epithelium proteins (KRT4, KRT13, SPRR3) showed consistent trends toward downregulation in PM patients compared to both healthy controls and lung cancer patients. This pattern might reflect a biological signature but it might also reflect altered respiratory mechanics characteristic of restrictive lung diseases, in which pleural restriction reduces tidal volumes and alters breathing efficiency ^40^, potentially affecting the mechanical desquamation of oral and esophageal epithelial cells during breath collection. This finding highlights a fundamental challenge in EBC proteomics: disease-related changes in respiratory mechanics can systematically bias protein collection, confounding the distinction between true biological signatures and sampling artefacts.

The analytical challenges we encountered in EBC proteomics, high missing data rates, low protein identification numbers and technical interferences, parallel those faced in single-cell proteomics, where similar statistical and technical hurdles exist [35,36]. Recent advances in single-cell proteomics methodology, including improved sample preparation techniques, boosting detection strategies, use of more sensitive mass spectrometers and specialized imputation algorithms designed for sparse datasets, may offer solutions applicable to EBC proteomics [37]. These emerging approaches could potentially be adapted to enhance biomarker discovery in dilute biological matrices like EBC.

In contrast, breath analysis has shown remarkable success for volatile organic compounds (VOCs) [38]. Several studies have demonstrated >90% sensitivity and negative predictive value for mesothelioma detection using GC-MS analysis, when discriminating PM patients from asbestos-exposed subjects [9,39,40]. These results suggest that the analysis of breath may be better suited for VOC-based biomarker discovery rather than protein-based approaches, given the superior analytical performance and clinical accuracy achieved with breath volatomics in respiratory oncology applications [41–43].

## Contributors

Conceptualization, K.L., J.P.V.M., K.G.; methodology, E.S., K.L., J.P.V.M., E.F., A.A., K.G.; formal analysis, E.S., E.F., A.A., K.G. and K.L.; investigation, E.S.; data curation, E.S., A.A., and E.F.; writing—original draft preparation, E.S. and E.F.; writing—review and editing, E.S., E.F., A.A., K.G., J.P.V.M., J.R. and K.L.; visualization, E.S., A.A., and E.F.; supervision, K.G., E.F., J.P.V.M. and K.L.; funding acquisition, K.G., K.L. and J.P.V.M. All authors have read and agreed to the published version of the manuscript.

## Acknowledgements

We wish to thank all colleagues from the Thoracic Oncology department of the Antwerp University Hospital, our colleagues from the VIB proteomics core, as well as staff of the collaborating companies (Joris Van Cleemput, Caroline Jonckheere and Tom Richart). We thank Leander Meuris from the VIB-UGent Center for Biomedical Biotechnology for assistance with MSqRob. We thank Kathleen Zwijsen and Eline Janssens for their assistance with the sample collection. We would also like to thank the patients and participants for their valuable contributions.

## Data Sharing Statement

The data presented in this study are available in the PRIDE archive. Project accession: PXD046980

