## Supplemental material for "Exhaled breath condensate proteomics using amphipols improves protein detection but reveals statistical challenges in respiratory disease biomarker discovery"

#### Supplemental methods

##### Sample preparation

Amphipols-based protein concentration

One mL of EBC was supplemented with 15 mM dithiothreitol, 150 mM NaCl and 50 mM sodium phosphate buffer ($NaPO_4$; pH 8) (figure 1a). Samples were heated for 5 min at 95 °C and immediately cooled down on ice for 5 min. Proteins were alkylated with 30 mM iodoacetamide and the pH was adjusted to 7.4-7.9 with 1 M NaOH. The reaction mixture was incubated for 30 min at room temperature in the dark. Amphipols A8-A35 (APols) were added to a final concentration of 1 mg/mL and samples were incubated at room temperature for 10 min. To precipitate the proteins, the samples were acidified with 10% formic acid to reach a pH of 3.3-3.5 and centrifuged for 7 min at 16,000 g at room temperature. After centrifugation, the supernatant was vacuum-dried and stored at -80 °C until further analysis and the pellet, containing the proteins, was washed with 125 µL of 0.01% formic acid and centrifuged for 3 min at 16,000 g. The washing solution was removed and the pellet was subsequently dissolved in 180 µL 30 mM triethylammonium bicarbonate (TEAB). Both endoproteinase Lys-C (100 ng/sample) and trypsin (200 ng/sample) were added simultaneously to the re-dissolved APols protein pellet and to the re-dissolved vacuum-dried supernatant resulting from APols precipitation. Samples were incubated overnight at 37 °C. The next day, the samples were acidified with 10% formic acid to reach a pH of 3.2-3.5 and centrifuged for 7 min at 16,000 g at room temperature. The supernatant, containing the tryptic peptides, was transferred to a new tube and stored at -80 °C until further analysis. Protein LoBind tubes were used to minimize protein loss. Samples were analyzed using LC-MS/MS in data-dependent (EBC replicates) or data-independent acquisition mode. Note that as a first test, the reduction/alkylation steps and the addition of endoproteinase Lys-C were not applied to the replicate EBC samples, similarly to the "Lyophilization method for protein concentration".

Lyophilization method for protein concentration

Four replicate EBC samples (two of 1 mL and two of 2 mL) were lyophilized. After lyophilization, the samples were resuspended in 100 µL of 10 mM TEAB pH 8.5. Proteins were denatured by placing the samples at 95°C for 5 min and immediately cooled down on ice afterwards. After adding 200 ng of trypsin, the samples were incubated overnight at 37°C. After digestion, the samples were acidified using 10% trifluoroacetic acid to reach a pH of 3.0 and then centrifuged for 3 min at 16,000g. The supernatant was transferred to a new protein LoBind tube and stored at -80 °C until further analysis. Samples were analyzed using LC-MS/MS in data-dependent acquisition mode.

#### Mass spectrometry analysis

Data-dependent acquisition

Peptides were desalted with OMIX C-18 tips (Agilent Technologies), dried and solubilized in 20 µL loading solvent A (0.1% TFA in water:acetonitrile (ACN) (98:2, v:v)) containing 2 mM TCEP before analysis. 15 µL of the sample was injected for LC-MS/MS analysis on an Ultimate 3000 RSLCnano system in-line connected to an Orbitrap Fusion Lumos mass spectrometer (Thermo). Trapping was performed at 10 µL/min for 4 min in loading solvent A on a 20 mm trapping column (made in-house, 100 µm internal diameter (I.D.), 5 µm beads, C18 Reprosil-HD, Dr. Maisch, Germany). The peptides were separated on a 200 cm µPAC™ column (C18-endcapped functionality, 300 µm wide channels, 5 µm porous-shell pillars, inter pillar distance of 2.5 µm and a depth of 20 µm; Thermo Fisher Scientific). The column was kept at a constant temperature of 50°C. Peptides were eluted by a linear gradient reaching 33% MS solvent B (0.1% formic acid (FA) in water:ACN (2:8, v/v)) after 105 min, 55% MS solvent B after 145 min and 99% MS solvent B at 150 min, followed by a 10-minutes wash at 99% MS solvent B and re-equilibration with MS solvent A (0.1% FA in water). The first 15 min, the flow rate was set to 750 nl/min after which it was kept constant at 300 nL/min.

The mass spectrometer was operated in data-dependent mode, automatically switching between MS and MS/MS acquisition. Full-scan MS spectra (300-1,500 m/z) were acquired in 3 s acquisition cycles at a resolution of 120,000 in the Orbitrap analyser after accumulation to a target AGC value of 200,000 with a maximum injection time of 30 ms. The precursor ions were filtered for charge states (2-7 required), dynamic range (60 s; ± 10 ppm window) and intensity (minimal intensity of 3E4). The precursor ions were selected in the multipole with an isolation window of 1.2 Da and accumulated to an AGC target of 5E3 or a maximum injection time of 40 ms and activated using HCD fragmentation (34% NCE). The fragments were analyzed in the Ion Trap Analyzer at normal scan rate.

Raw spectra files were searched with the MaxQuant software (v 2.0.3.0) with the Andromeda search engine at default search settings including an FDR set at 1% on both the peptide and protein level. Files from both experimental groups, APols and lyophilization, were searched independently. Spectra were searched against the reviewed human UniProt database (January 2023). The mass tolerance for precursor and fragment ions was set to 20 ppm and 4.5 ppm, respectively, during the main search. The enzyme specificity was set to trypsin, with a maximum of two missed cleavages allowed. Variable modifications were set to oxidation (Met) and protein N-terminal acetylation. A minimum of one peptide was required for identification. Proteins were quantified by the MaxLFQ algorithm integrated in the software and a minimum ratio count of two unique or razor peptides was required for quantification.

Data-independent acquisition

Samples were run in data-independent mode on an Evosep One LC-system (Evosep, Denmark) in-line connected to a Q Exactive HF mass spectrometer (Thermo). Peptides were analyzed with the Whisper 20 SPD method using the endurance Evosep column (15 cm x 75 µm I.D., 1.9 µm beads, EV-1112, Evosep, Denmark) connected to a fused silica emitter (10 µm inner diameter) (EV-1111, Evosep, Denmark). For elution of the peptides from the column, 0.1% FA in LC-MS-grade water and 0.1% FA in ACN were used as mobile phases. Full-scan MS spectra ranging from 375-1,500 m/z with an AGC target value of 5E6, a maximum fill time of 50 ms and a resolution at 200 m/z of 60,000 were followed by 30 quadrupole isolations with a precursor isolation width of 10 m/z for HCD fragmentation at an NCE of 30% after filling the trap at a target value of 3E6 for maximum injection time of 45 ms. MS2 spectra were acquired at a resolution of 15,000 at 200 m/z in the Orbitrap analyser without multiplexing. The isolation intervals ranging from 400 – 900 m/z were created with the Skyline software tool. QCloud was used to control instrument longitudinal performance during the project [1].

Data-independent acquisition spectra were searched with the DIA-NN software (v1.8.1) in library-free mode against the UniProt database (May 2021 release). The mass accuracy was set to 20 ppm and the MS1 accuracy to 10 ppm, with a precursor FDR of 0.01. Enzyme specificity was set to trypsin/P with a maximum of two missed cleavages. Variable modifications were set to oxidation of methionine residues (to sulfoxides) and acetylation of protein N-termini. Carbamidomethylation of cysteines was set as a fixed modification. The peptide length range was set to 7-30 residues with a precursor charge state between 1-4. The m/z range was set between 400-900 and 200-1,800 for the precursor and fragment ions, respectively. Cross-run normalization was set to RT dependent with the quantification strategy set to high accuracy and the neural network classifier to double-pass mode. The result report file was further processed for statistical analysis.

#### Statistical analysis

For the analysis of APols recovery and the comparison with the lyophilization protocol, one replicate of the APols group was excluded due to sample loss during the sample preparation. Peptide and protein intensities were obtained from the peptides.txt and proteinGroup.txt files generated by MaxQuant from analysis of APols precipitates, supernatants (SN) and lyophilized samples. Only proteins and peptides mapped against *H. sapiens* were retained. Proteins and peptides identified only by site or marked as reverse database hits were filtered prior further analysis. Percentage of recovery efficiency was calculated with peptides only identified in both fractions: APols precipitates and SN. Recovery efficiency for shared peptides was calculated as follows: for each peptide sequence, the total intensity was calculated as the sum of intensities from the APols precipitate and SN fractions. Recovery efficiency (%) was then calculated as:

$$\text{Recovery Efficiency (\%)} = (\text{Intensity}_{\text{APols}}/(\text{Intensity}_{\text{APols}} + \text{Intensity}_{\text{SN}})) \times 100$$

The median recovery efficiency and interquartile range (IQR) were calculated across all shared peptides. Peptides with recovery efficiency greater than 90% and 95% were quantified to assess the proportion of highly efficiently recovered peptides. Coefficients of variation (CV) were calculated based on the sample median-normalized raw peptide intensities and on the label-free quantification (LFQ) intensities of proteins. CVs were calculated from peptides or proteins that were consistently identified in at least three replicates within both APols and lyophilization groups. Statistical analysis was performed using a paired Wilcoxon signed-rank test.

Total quantity of precursor intensities was obtained from the DIA-NN stats file. Precursor intensities were obtained from the DIA-NN report and imported into a QFeatures [2] object at the precursor level, using precursor quantity as the quantitative measure. Precursors were retained only if they were proteotypic and had a q-value < 0.01, and proteins were considered only if supported by more than two peptides. To further reduce noise, precursors were required to have at least 30% valid values in at least one clinical sample group, resulting in a total of 2,374 precursors. Precursor intensities were $log_2$-transformed and normalized across samples using median centering.

Batch correction and missing-value imputation were performed with omicsGMF [3], starting from normalized precursor intensities and correcting for batch and device effects. Imputed precursor intensities were aggregated to the protein level using median polish summarization, yielding 248 proteins. Protein-level differential expression analysis was performed with a linear model implemented in MSqRob [3–5], adjusting for all available cohort-specific covariates (smoking status, pack-years, gender, BMI, batch and device). False discovery rate (FDR)-adjusted p-values were used to correct for multiple testing. Statistical significance was defined as an adjusted p-value < 0.05.

The tissue category annotation from the Human Protein Atlas (v24.0) was consulted to annotate tissue-enriched proteins [6]. Supervised and unsupervised statistical analyses (Permutational Multivariate Analysis of Variance, Analysis of Similarities, calculation of average silhouette width and the Adjusted Rand Index) were calculated with the Vegan package from RStudio. Plots and Figures were generated in RStudio.

#### Supplemental figure and table legends

**Supplemental figure 1.** APols precipitation efficiency. Recovery efficiency was calculated as the ratio of peptide intensity in APols precipitate to total intensity (APols + supernatant) × 100. The cumulative distribution represents the percentage of peptides achieving at least the indicated recovery efficiency. Dashed lines: 90% and 95% recovery thresholds, n = 362 peptides.

**Supplemental figure 2.** Fold change ($log_2$) distribution of shared peptide intensities between APols and lyophilization sample preparation methods. Each dot represents a shared peptide, with blue dots indicating higher intensity using the APols method and grey dots indicating higher intensity using lyophilization. The boxplot shows the median fold change and interquartile range in $log_2$ units. The half-violin plot illustrates the overall distribution. The horizontal dashed line at zero represents equal intensity. $Log_2$ of median-fold change = 1.11, paired Wilcoxon signed-rank test, $p = 1.11e-55$, n = 554 shared peptides.

**Supplemental figure 3.** Precursor completeness. 2,374 unique precursors (x-axis) ranked by completeness (y-axis) across the 145 EBC samples. Dashed red line: 346 precursors account for 100% completeness. Dashed blue line: 947 precursors account for 80% completeness.

**Supplemental figure 4.** Distribution of the intensities of 2,374 precursors (x-axis) collected with the TurboDECCS (a) or EcoScreen (b) devices, plotted against percentage of completeness (y-axis). Similar number of precursors are binned (n = 79-80). Intensities are normalized and $log_2$ transformed. Boxes show the interquartile range and the median. Red dots show the averages of the completeness percentage.

**Supplemental figure 5.** Box plots showing the coefficient of variation (CV) distribution for 2,374 precursors corrected with omicsGMF by batch and device effect measured across the four clinical groups. Each box represents the interquartile range ($25^{th}$-$75^{th}$ percentile), with the median indicated by the central line. Whiskers extend to the most extreme data points within 1.5 times the interquartile range. AEX: asbestos-exposed, HC: healthy controls, LC: lung cancer, and PM: pleural mesothelioma.

**Supplemental figure 6.** Principal Component Analysis of 145 EBC samples after batch and device effect correction. Scatter plot showing the distribution of samples along the first two principal components (PC1 and PC2) based on protein expression profiles. Samples are coloured by collection device (purple: TurboDECS, green: EcoScreen) and shaped by processing batch (circles: Batch1, triangles: Batch2, squares: Batch3, crosses: Batch4, diamonds: Batch5). PC1 = 23.31%, PC2 = 15.99%, PC3 = 15.05%, PC4 = 8.08.

**Supplemental table 1.** Peptides identified in the APols precipitates (Apols), supernatants (SN) and lyophilized (Lyophil) samples across replicates. Peptide sequence, raw intensity and protein accession (UniProt, version January 2023) are indicated.

**Supplemental table 2.** ProteinGroups file obtained from MaxQuant (v 2.0.3.0). Majority protein IDs, protein IDs, group (Apols, Lyophil and SN), replicate number and LFQ intensity are shown.

**Supplemental table 3.** 362 peptides commonly identified in APols precipitates and supernatants (SN). Intensities are calculated as the sum of the intensities across replicates. Recovery percentage is calculated as the intensity of APols percentage versus the total intensity (APols + SN) for each peptide. The fold enrichment is calculated as the intensities of APols versus SN.

**Supplemental table 4.** Total intensity of the APols and supernatants (SN) replicates. The fold enrichment is calculated as the intensities of APols versus SN.

**Supplemental table 5.** 24 tissue-enriched proteins annotated in the Human Protein Atlas (v24.0). Majority protein IDs, group (APols and Lyophilization), LFQ intensity, replicate and tissue annotation are indicated.

**Supplemental table 6.** Tissue precursor annotation based on the Human Protein Atlas (HPA) (v24.0). Individual normalized precursor intensities were summarized by tissue in each sample. Percentage of enriched tissue was calculated as intensity tissue / total intensity. Intensity from non-annotated precursors was include to compute the total precursor intensity.

**Supplemental table 7.** 248 protein intensities across 145 EBC samples. Batch, group, age, BMI, gender, smoking status, device, packyears and protein accessions (UniProt) are indicated. Protein intensities result from precursor normalization and aggregation, performed in MSqRob.

**Supplemental table 8.** Pairwise PERMANOVA comparisons between clinical groups. F_model (Pseudo-F): Pseudo-F statistic from PERMANOVA, measuring the ratio of between-group to within-group multivariate distances. $R^2$: Coefficient of determination, representing the proportion of multivariate variance explained by the grouping factor (ranges from 0 to 1). Pr_F (*p*-value): Permutation-based *p*-value indicating the probability of observing the F-statistic under the null hypothesis of no difference in multivariate composition between groups (values < 0.05 are considered statistically significant; based on permutation tests).

**Supplemental tables 9-16.** Results of the pairwise linear model comparisons across the indicated contrast.

Supplemental figure 1

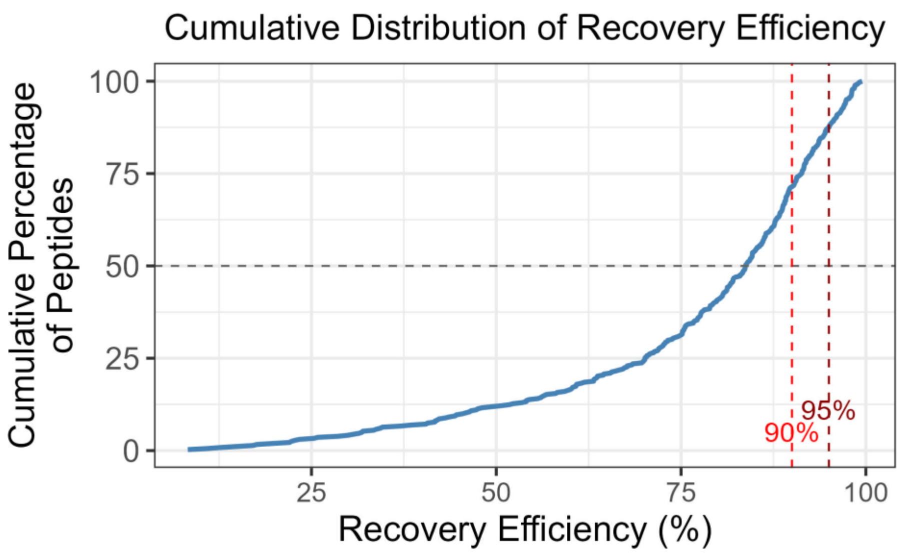

Supplemental figure 2

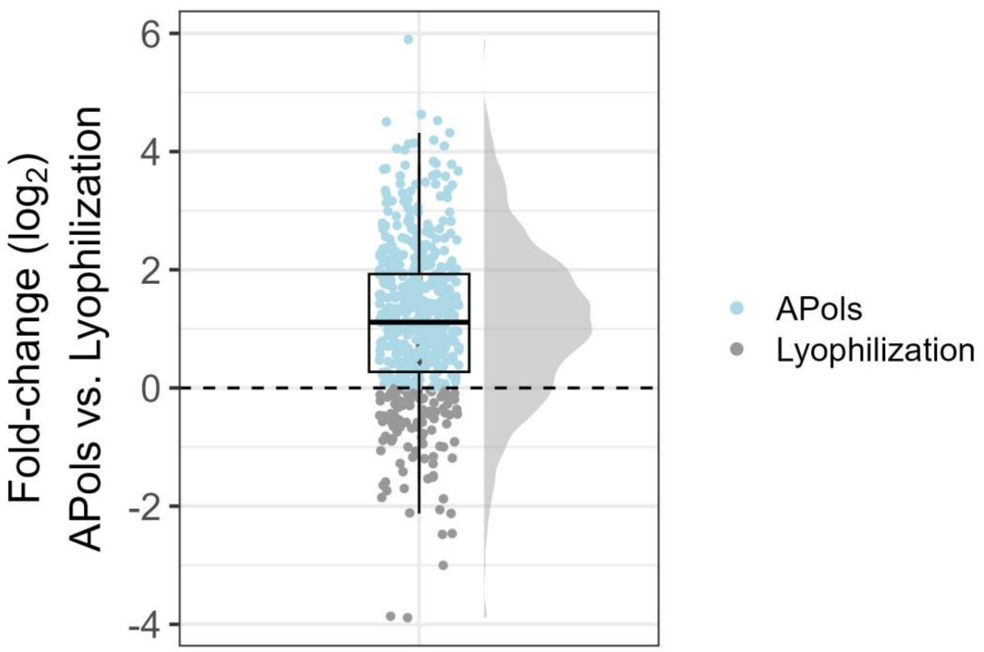

Supplemental figure 3

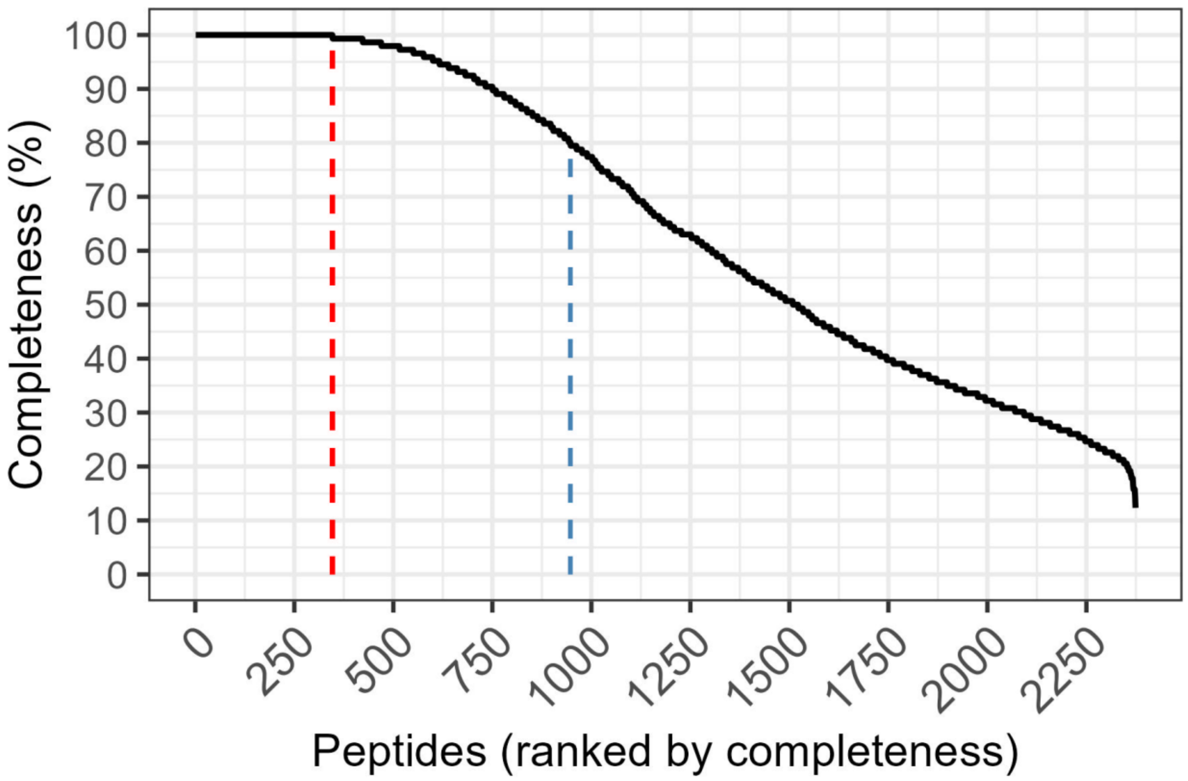

### Supplemental figure 4

#### a) TurboDECCS

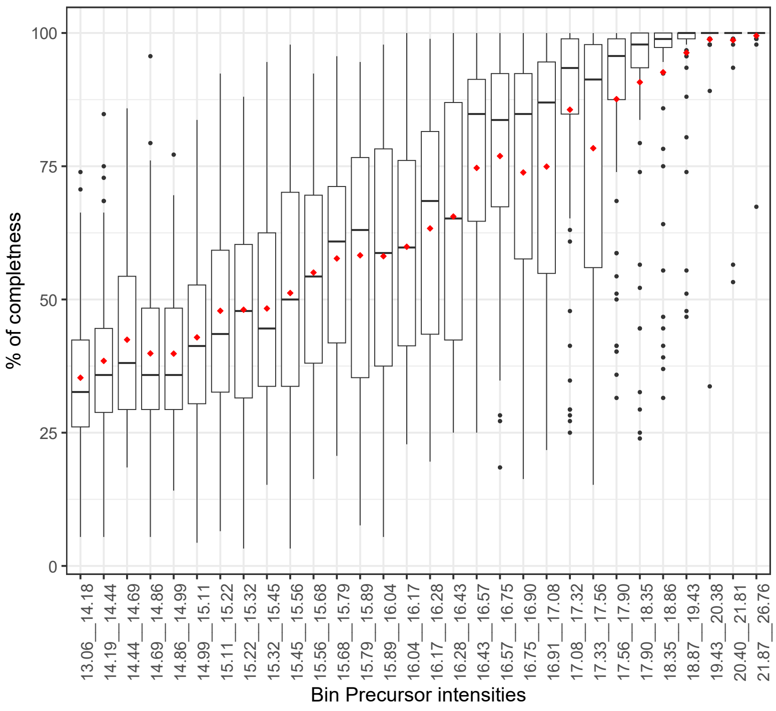


#### b) EcoScreen

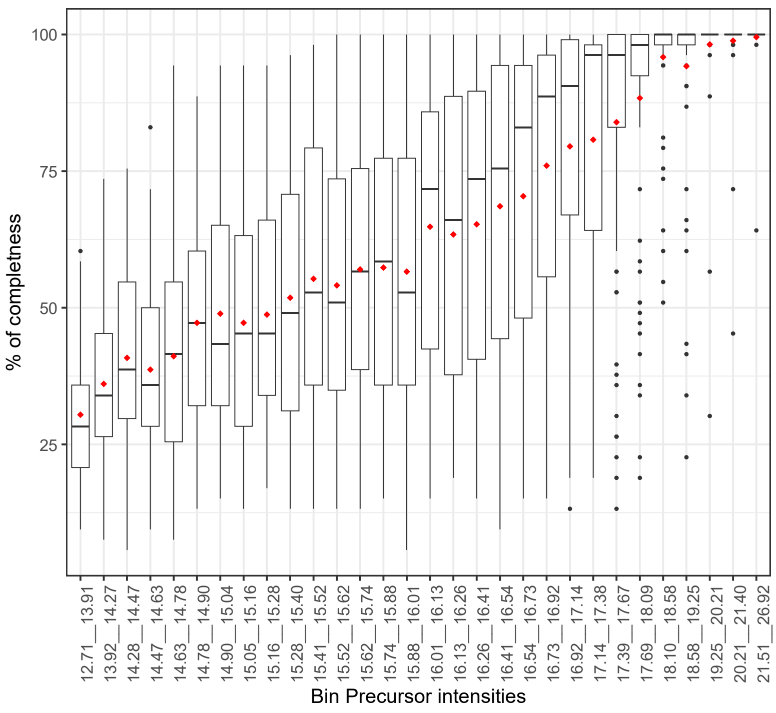

Supplemental figure 5

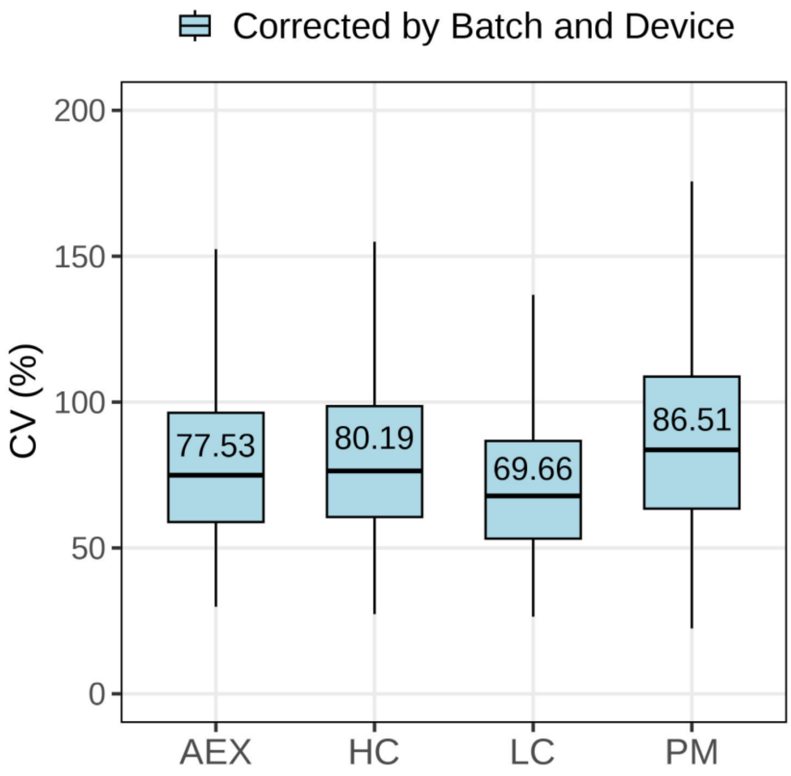

Supplemental figure 6

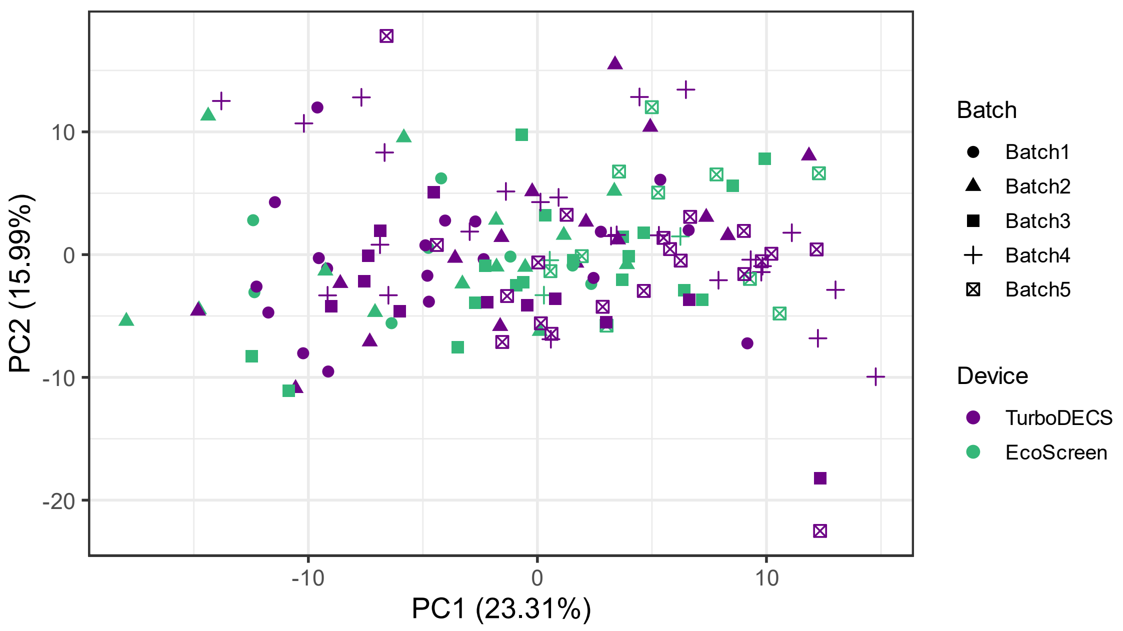
